# Extracellular vesicle-mediated suppression of macrophage STING signaling promotes immune dysfunction in dedifferentiated liposarcoma

**DOI:** 10.64898/2026.08.07.743624

**Authors:** Qi Zhang, Jay K Mandula, Patricia Sarchet, Priya Dhawale, Fernanda C. C. de Faria, Timothy Zhang, Sydney Rentsch, Premanshu K Singh, Ali F Usmani, Roma Karna, Connor P Harper, Valerie Grignol, Jing Wang, Yong Zhang, Zihai Li, Raphael E Pollock, Federica Calore

**Affiliations:** Department of Surgery, Division of Surgical Oncology, The James Comprehensive Cancer Center, The Ohio State University Wexner Medical Center, Columbus, OH, USA; Department of Retroperitoneal and Soft Tissue Surgical Oncology, Zhongshan Hospital, Fudan University; Pelotonia Institute for Immuno-Oncology, The Ohio State University Comprehensive Cancer Center-James Cancer Center and Solove Research Institute; The Ohio State University College of Arts and Sciences; Department of Mechanical and Aerospace Engineering, The Ohio State University; Department of Cancer Biology and Genetics, The Ohio State University

**Keywords:** Dedifferentiated liposarcoma, extracellular vesicles, macrophage polarization, stimulator of interferon genes (STING) signaling, innate immunity, tumor microenvironment

## Abstract

**Background:** Dedifferentiated liposarcoma (DDLPS) is characterized by abundant immune cell infiltration yet derives limited benefit from immune checkpoint blockade and stimulator of interferon genes (STING) agonist-based strategies, suggesting tumor-mediated suppression of antitumor immunity. Tumor-associated macrophages are the most abundant immune populations in DDLPS, but the factors regulating their function remain incompletely understood.

**Methods:** Extracellular vesicles (EVs) were isolated from two DDLPS cell lines and serum from 16 DDLPS patients and 13 healthy donors. EVs’ impact on cyclic guanosine monophosphate-adenosine monophosphate (cGAMP) -induced macrophage activation was assessed by cytokine secretion, surface markers, functional assays and macrophage-T-cell coculture. Proteomics was performed in EV-treated and EV-untreated macrophages from three donors. Pathway and protein interaction analyses were integrated with The Cancer Genome Atlas (TCGA) DDLPS transcriptomic and survival data.

**Results:** We show that EVs released by DDLPS cells suppress macrophage responsiveness to classic STING agonist cGAMP. EVs derived from DDLPS attenuated cGAMP-induced expression of type I interferon-associated cytokines and chemokines, reduced IFN-β secretion, and impaired phosphorylation of STING, TBK1 and IRF3. Functionally, DDLPS EV exposure shifted macrophages toward an immunoregulatory phenotype, restrained phagocytic activity, and attenuated macrophage-dependent T-cell proliferation while promoting T-cell exhaustion. Proteomic profiling revealed extensive macrophage reprogramming characterized by suppression of STING-associated signaling, antigen processing and presentation associated pathways and proteins targeted by miR-16-5p. Consistent with these findings, STING expression was associated with prolonged overall survival in DDLPS, while reduced expression of miR-16-5p target proteins was associated with attenuated STING pathway activity and immunostimulatory macrophage signatures.

**Conclusions:** These findings identify EV-mediated suppression of macrophage STING signaling as a mechanism of immune dysfunction in DDLPS and provide a framework for understanding immune resistance in this disease.

## Introduction

Retroperitoneal dedifferentiated liposarcoma (DDLPS) is one of the most clinically challenging soft-tissue sarcomas and is particularly problematic because of its deep anatomic location, large tumor burden at presentation, limited systemic treatment options and poor prognosis(1–3). Complete surgical resection remains the cornerstone of therapy for localized disease, but the major clinical challenge in retroperitoneal DDLPS is repeated locoregional recurrence rather than distant metastasis. Even following compartmental resection as recommended by the transatlantic Australasian retroperitoneal sarcoma working group (TARPSWG), recurrence rates remain high, and repeated operations are often associated with progressive loss of organ function and diminishing opportunities for further curative surgery(4, 5). Several clinical trials have shown limited efficacy of immune checkpoint blockade (ICB) in DDLPS(6). Similarly, early clinical studies investigating stimulator of interferon genes 1 (STING) agonists have reported substantially lower response rates in liposarcoma compared with other solid tumors(7). Together, these observations suggest the existence of tumor-intrinsic mechanisms that suppress antitumor immunity and limit STING pathway activation in DDLPS.

STING is a critical player in the innate and adaptive immune response. Upon detection of cytosolic double stranded DNA or noncanonical activators, cyclic GMP-AMP synthase (cGAS) catalyzes production of cyclic guanosine monophosphate-adenosine monophosphate (cGAMP) from adenosine and guanosine triphosphate (ATP, GTP). Binding of STING to cGAMP triggers recruitment and activating phosphorylation of STING-complex members such as TANK-binding kinase 1 (TBK1) which subsequently coordinate phosphorylation of interferon response factor 3 (IRF3), resulting in phospho-IRF3 dimerization and nuclear translocation. In the nucleus, phospho-IRF3 binds to interferon stimulated response elements (ISREs) in the promoters of target genes, inducing expression of ISRE-regulated genes such as type I interferons. Importantly, in the tumor microenvironment, cGAS-STING pathway induced production of type I interferon signaling promotes immunostimulatory macrophage polarization and macrophage-directed T cell activation and antitumor immune responses(8). Despite growing therapeutic interest in STING agonists, the mechanisms responsible for impaired STING signaling in liposarcoma remain poorly understood.

As a bridge between innate immunity and adaptive immunity, macrophages are the most abundant immune population of the tumor microenvironment (TME), accounting for approximately 50% of tumor-infiltrating hematopoietic cells(9). Prior reports have demonstrated that immune cell infiltration predicts clinical outcome in DDLPS, with accumulation of alternatively activated “M2” macrophage associated with worse outcomes(10). These findings suggest that macrophage phenotype may play an important determinant of immune suppression and invasion within the DDLPS tumor microenvironment. However, the mechanisms by which liposarcoma reprograms macrophage function and impairs antitumor immunity remain incompletely defined.

Extracellular vesicles (EVs) have emerged as important mediators of tumor-host communication and can regulate innate immune signaling, macrophage polarization, and antitumor immunity (11). Given the role of EVs as regulators of macrophage function, we hypothesized that DDLPS-derived EVs may contribute to suppression of STING-mediated antitumor responses. Our previous studies have shown that DDLPS-derived EVs carrying *MDM2* DNA promote pro-tumorigenic reprogramming of preadipocytes, while EV-associated microRNAs (miRNA) miR-25-3p and miR-92a-3p stimulate macrophage IL-6 secretion and promote liposarcoma progression(12, 13). Moreover, additional studies have indicated that EV-associated miR-16-5p impacts macrophage polarization resulting in repression of immunostimulatory “M1” phenotypes while additional miRNAs have been reported to more broadly tune myeloid functions in cancer(14). However, the role of EV encapsulated miRNAs and their impact of macrophage phenotype, function and immune outcomes in DDLPS remain poorly characterized. Together, these observations indicate that DDLPS-derived EVs actively remodel the tumor microenvironment in ways that favor disease progression and suggest that EVs may influence macrophage-mediated immune responses through mechanisms extending beyond inflammatory cytokine production.

Here, we investigated whether DDLPS-derived EVs suppress macrophage-mediated antitumor immune activation through inhibition of the STING pathway. We demonstrate that DDLPS-derived EVs impair activation of the STING-TBK1-IRF3 signaling axis, suppress type I interferon responses, promote immunoregulatory macrophage polarization, and impair macrophage-mediated T-cell activation. Integrated proteomic and pathway analyses further identify a miR-16-5p-associated signature linked to reduced STING pathway activity and impaired immunostimulatory macrophage function.

## Results

### DDLPS-derived EVs suppress macrophage type I interferon responses following cGAMP stimulation

Previous studies have linked tumor-derived extracellular vesicles (EVs) to suppression of antitumor immune responses, including modulation of type I interferon signaling(15, 16). To investigate whether DDLPS-derived EVs regulate macrophage innate immune responses, we isolated EVs from the conditioned medium of DDLPS cell lines, Lipo141 and Lipo246, using ultracentrifugation. Isolated vesicles displayed characteristic morphology, size distribution, and protein marker expression consistent with MISEV guidelines(17) (Figure S1A-C).

Because activation of the STING pathway induces type I interferon responses that support antitumor immunity, we examined the effects of DDLPS-derived EVs on cGAMP-stimulated human macrophage cell lines. Prior studies have linked EV exposure to altered induction of antitumor immune responses including EV exposure-linked changes in expression of type I interferon(15, 16). Exposure to DDLPS-derived EVs significantly reduced interferon beta 1 (ifnb1) and interferon alpha 2 (ifna2) mRNA expression after cGAMP stimulation (Figure 1A). In addition, we observed decreased secretion of IFN-β protein after EV exposure (Figure 1B).

**Figure 1:**
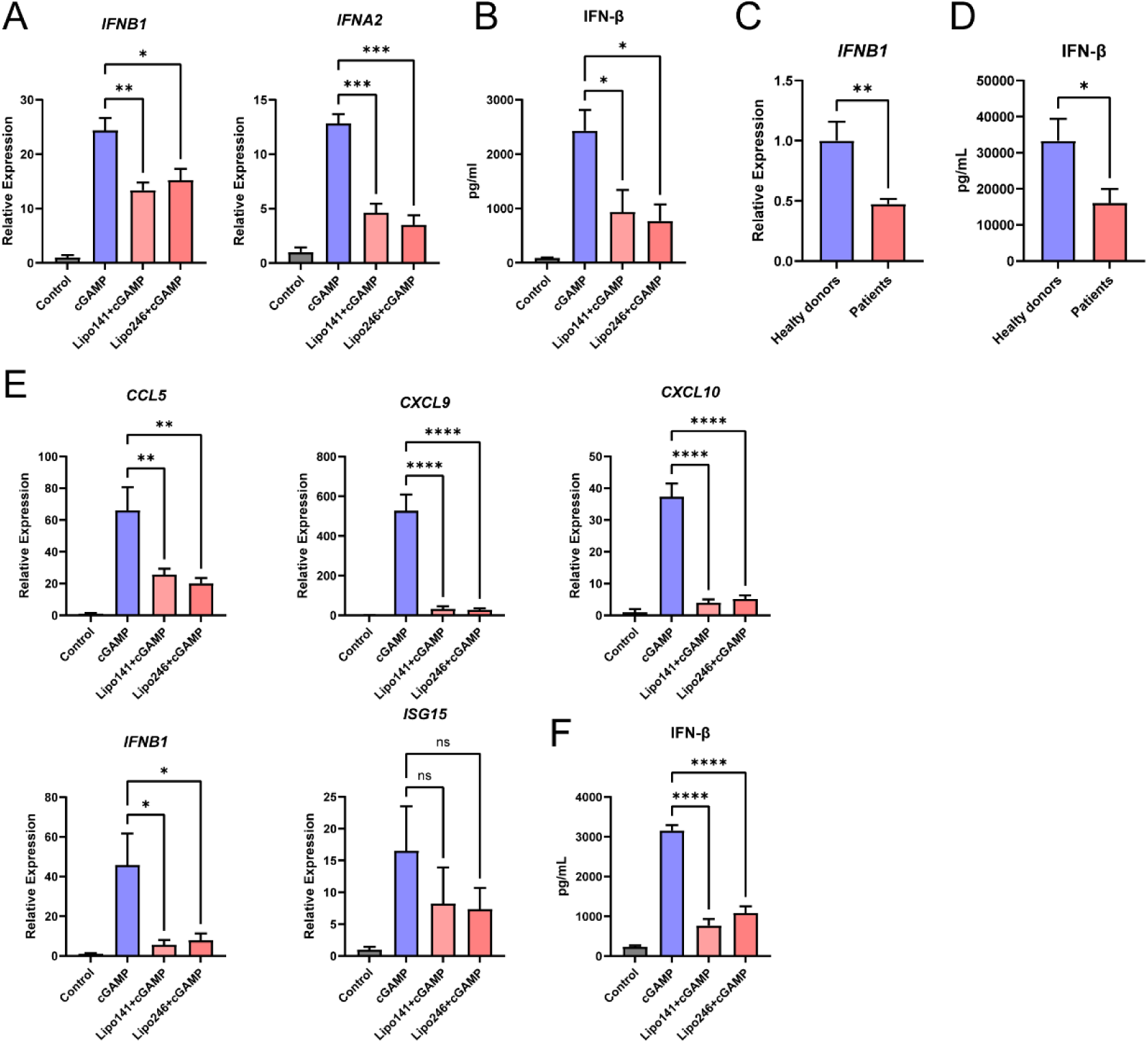
Exposure to DDLPS-EVs restricts cGAMP induced expression of cytokine in human macrophages: (A, C) mRNA expression of U937-derived macrophages incubated with EVs isolated from DDLPS cell lines (A) or human serum samples (C) (n=13 healthy donors, n=16 DDLPS patients) for 24 h, followed by cGAMP stimulation for 8h. (B, D) IFN-β protein secretion in culture supernatants from U937-derived macrophages exposed to EVs isolated from DDLPS cell lines (B) or human serum samples (D), assessed by ELISA following 24 h EV exposure and 8h cGAMP stimulation. (E) mRNA expression in human monocyte-derived macrophages (MDMs) incubated with EVs isolated from Lipo141 or Lipo246 for 24 h, followed by cGAMP stimulation for 8 h. (F) IFN-β secretion in culture supernatants from MDMs analyzed by ELISA following EV exposure and cGAMP stimulation.

To validate these findings in patient-derived samples, we isolated EVs from the serum of 16 DDLPS patients and 13 healthy donors. Nanoparticle Tracking Analysis (NTA) showed comparable vesicle size distributions between the two groups, although DDLPS patient serum contained slightly higher EV concentrations (Figure S1D). Consistent with the findings obtained using cell line-derived EVs, EVs isolated from DDLPS patient serum significantly reduced Ifnb1 mRNA expression (Figure 1C) and IFN-β protein secretion from U937-derived macrophages following cGAMP stimulation (Figure 1D).

To further characterize the impact of DDLPS EV exposure, we isolated CD14+ monocytes from healthy donor human peripheral blood and differentiated them into macrophages (Monocyte Derived Macrophages, MDM) *in vitro* prior to treatment with DDLPS EVs. Remarkably, DDLPS EVs exposure induced downregulation of *CCL5*, *CXCL9*, *CXCL10*, *IFNB1* and *ISG15* mRNA expressions in response to cGAMP stimulation (Figure 1E). In line with these transcriptional changes, DDLPS EVs significantly decreased the secretion of IFN-β protein by MDMs (Figure 1F).

Collectively, these findings demonstrate that DDLPS-derived EVs suppress macrophage type I interferon responses following STING activation.

### DDLPS-derived EVs promote immunoregulatory polarization of macrophages

STING activation promotes immunostimulatory macrophage polarization, with induction of an immunostimulatory “M1” macrophage phenotype, culminating in upregulation of antigen processing/presentation and expression of costimulatory markers(18). We therefore investigated whether DDLPS-derived EVs induced this effect. cGAMP stimulation increased expression of M1-associated markers in MDMs, including Human Leukocyte Antigen – DR isotype (HLA-DR) and cluster of differentiation 86 (CD86), whereas cotreatment with DDLPS-derived EVs and cGAMP impaired upregulation of these markers (Figure 2A-B, E). Conversely, DDLPS-derived EVs increased expression of the immunosuppressive “M2” associated markers including programmed death ligand 1 (PD-L1) (19–21)and cluster of differentiation 206 (CD206) (Figure 2C-D, E).

**Figure 2:**
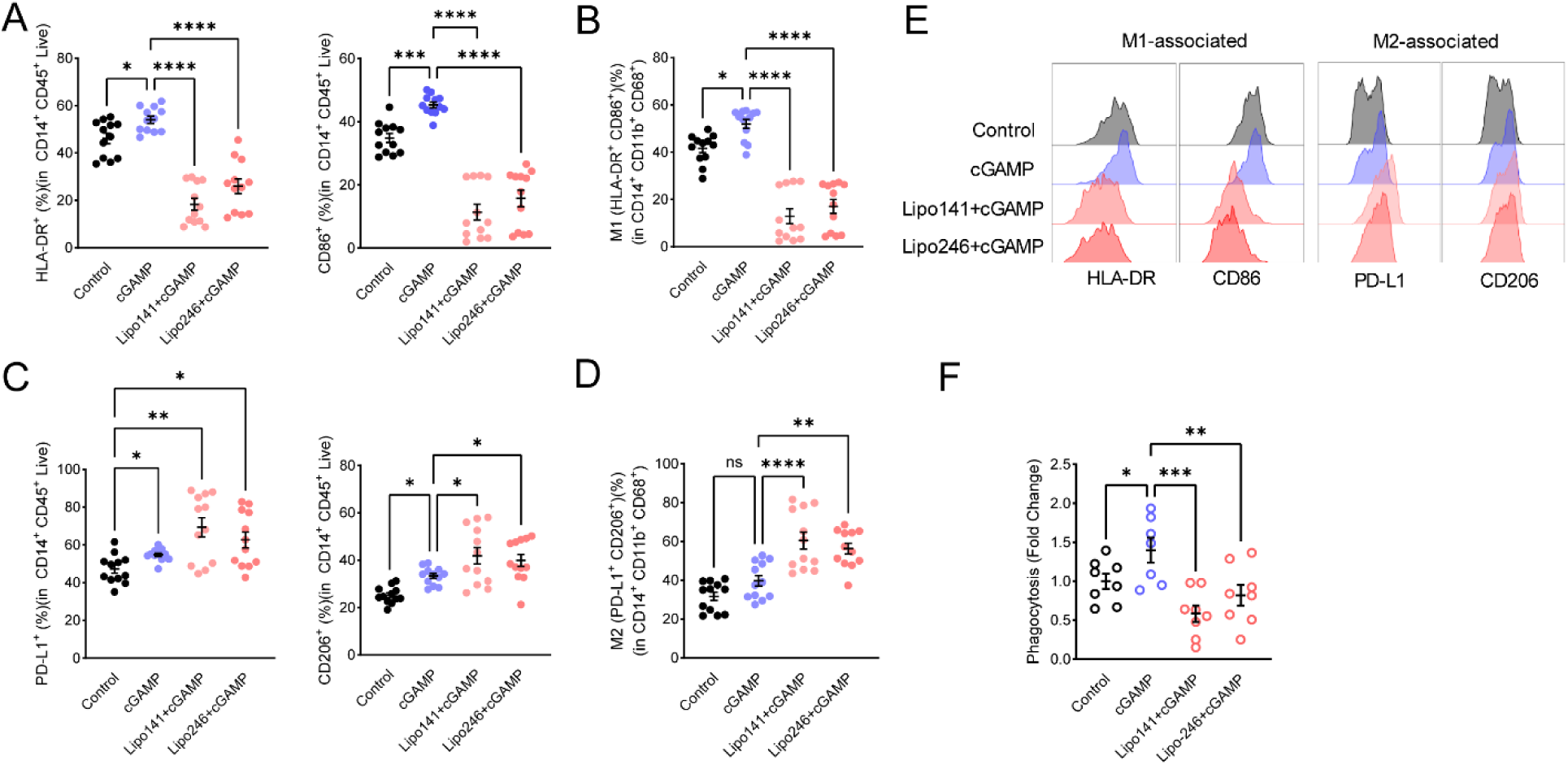
DDLPS-EV exposure promotes immunoregulatory macrophage polarization: (A) Flow cytometry analysis showing reduced expression of the M1-associated markers HLA-DR (left) and CD86 (right) in macrophages treated with DDLPS-derived EVs following cGAMP stimulation. (B) Quantification of M1-polarized macrophages following EV exposure and cGAMP stimulation. (C) Flow cytometry analysis showing increased expression of the immunosuppressive markers PD-L1 (left) and CD206 (right) in macrophages treated exposed to DDLPS-derived EVs. (D) Quantification of M2-polarized macrophages following EV exposure and cGAMP stimulation. (E) Representative histograms showing expression of M1-associated (HLA-DR, CD86) and M2-associated (PD-L1, CD206) markers in macrophages. (F) Phagocytosis assay showing reduced uptake of fluorescently labeled latex beads in macrophages exposed to DDLPS-derived EVs following cGAMP stimulation.

Immunostimulatory polarization promotes macrophage phagocytic activity, with phagocytosed material supplying antigen for subsequent presentation. We therefore investigated whether DDLPS-derived EV exposure altered macrophage phagocytic function. Markedly, while cGAMP treatment increased phagocytic uptake of fluorescently labeled beads, exposure to DDLPS EVs impaired macrophage phagocytosis (Figure 2F).

Since immunostimulatory macrophages support adaptive immune responses through T cell expansion and proliferation via antigen presentation and expression of costimulatory molecules, we next asked whether EV-mediated macrophage reprogramming altered T-cell activation. MDMs were treated with vehicle, cGAMP alone, or cGAMP together with DDLPS-derived EVs prior to coculture with matched human donor T cells for 72 hours. Compared with CD4+ or CD8+ T cells cocultured with cGAMP-stimulated MDMs, T cells exposed to DDLPS EV-treated MDMs exhibited reduced proliferation (Figure 3A). Consistent with induction of T cell dysfunction, exposure to DDLPS EV-treated MDMs resulted in increased expression of exhaustion markers programmed cell death protein 1 (PD1) and T cell immunoglobulin and mucin domain-containing protein 3 (TIM3) in CD4+ and CD8+T cells (Figure 3B).

**Figure 3:**
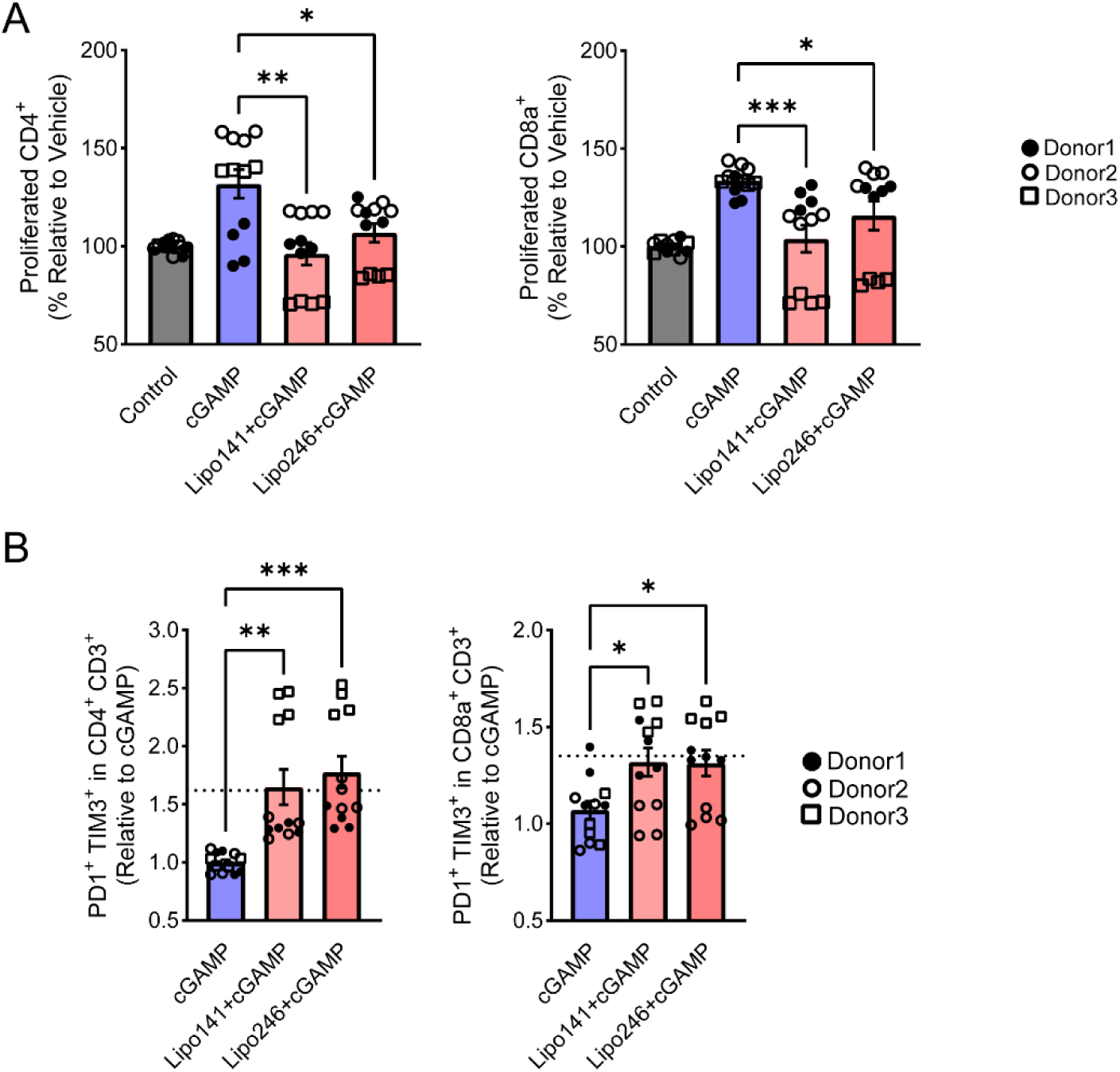
Conditioning with DDLPS-EVs restricts macrophage induced T cell proliferation and promotes exhaustion: (A) Flow cytometric analysis of CD4 (*left*) and CD8 (*right)* T cell proliferation after 72 hours coculture with vehicle, cGAMP, EV141+cGAMP or EV246+cGAMP treated MDM from matched donors. (B) Fold change in the levels of terminally exhausted (TIM3^+^ PD1^+^) CD8 (*left)* or CD8 (*right)* T cells in total live T cells after 48 hours coculture with vehicle, cGAMP, EV141+cGAMP or EV246+cGAMP treated MDM from matched donors.

Together, these findings indicate that DDLPS-derived EVs reprogram macrophages towards an immunoregulatory phenotype characterized by impaired innate and adaptive immune stimulatory functions.

### DDLPS-derived EVs inhibit STING pathway activation in macrophages

Since DDLPS-derived EVs suppressed macrophage type I interferon responses following stimulation with the canonical STING agonist cGAMP, we next investigated whether EV exposure directly altered STING pathway activation. Kinetic evaluation of STING signaling in MDMs demonstrated that DDLPS-derived EVs reduced STING phosphorylation as early as 1 h after cGAMP stimulation (Figure 4A). Furthermore, DDLPS EV exposure decreased IRF3 phosphorylation at 1 h after cGAMP treatment, with maximal inhibition observed 3h following cGAMP stimulation (Figure 4A). Consistent with impaired STING pathway activation, TBK1 phosphorylation was likewise reduced in DDLPS EV-macrophages (Figure 4A). Notably, IRF3 has been previously reported to regulate expression of CD86 and HLA-DR(22), indicating that the decreased expression of CD86 and HLA-DR apparent in DDLPS EV treated macrophages may occur due to attenuation of STING driven IRF3 activation.

**Figure 4:**
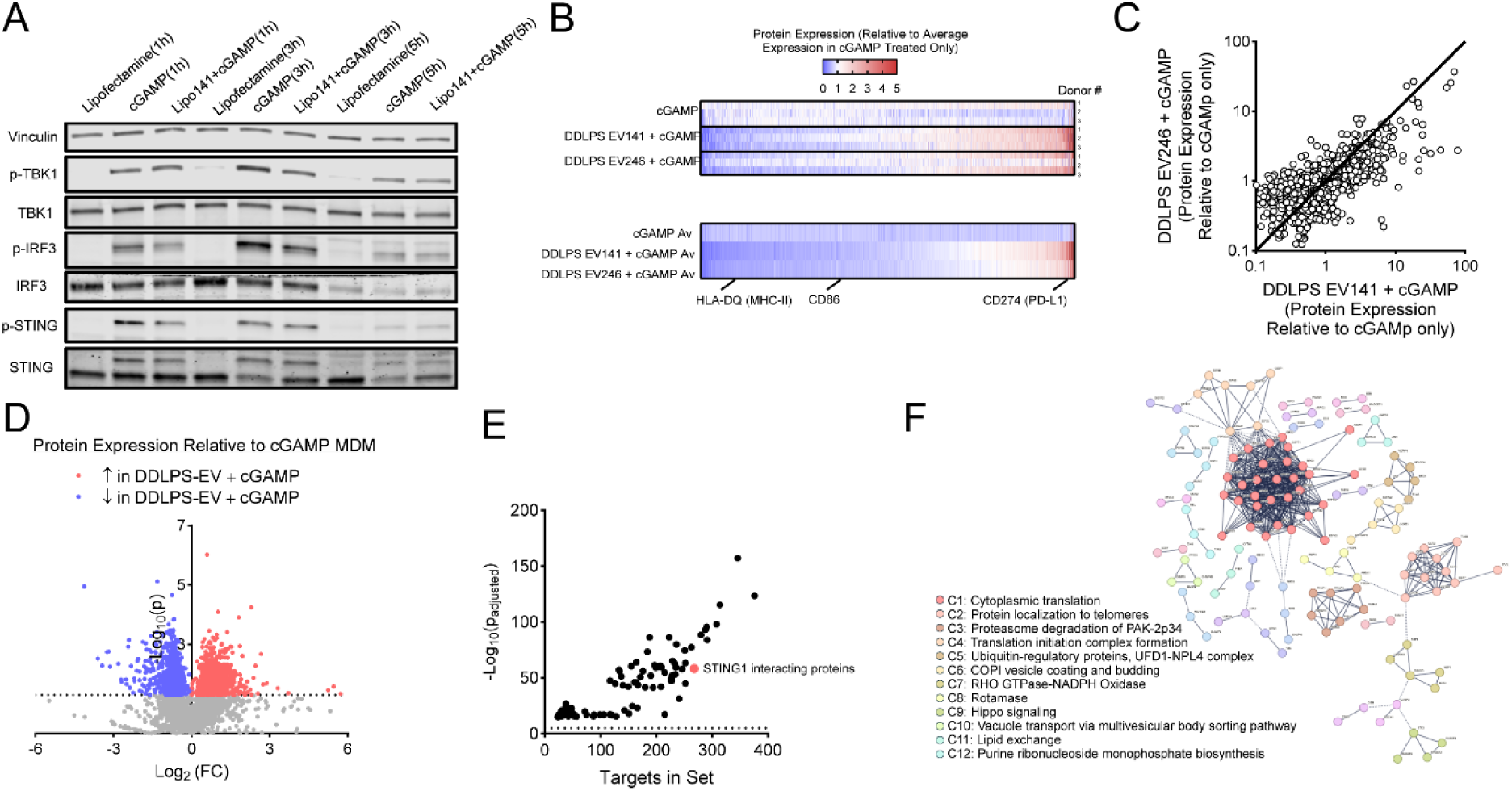
DDLPS-EVs impair STING pathway activation in response to cGAMP stimulation: (A) Western blot of the levels of phosphorylated proteins (STING, TBK1, and IRF3) in MDMs incubated with DDLPS-derived EVs for 24 h, followed by cGAMP stimulation. (B) Heat map visualization for individual sample protein (*top*) abundance (MS2 normalized) expression as value/average expression in cGAMP treated MDM only across all donors and heat map visualization (*bottom*) for average protein abundance (MS2 normalized) expression as value/average expression in cGAMP treated MDM only across all donors with highlighted proteins of interest HLA-DQ (MHC-II), CD86 and CD274 (PD-L1). (C) Expression correlation comparison between MDM treated with DDLPS EV141 + cGAMP versus DDLPS EV246 + cGAMP relative to cGAMP treated MDM only. (D) Volcano plot visualization for Log2 fold change of average protein expression in MDM samples treated with DDLPS EV141 or 246 + cGAMP relative to cGAMP with highlight for 1036 proteins significantly increased in MDM treated with DDLPS EV141 or 246 + cGAMP relative to cGAMP and 782 proteins significantly decreased in MDM treated with DDLPS EV141 or 246 + cGAMP relative to cGAMP. (E) Visualization of pathways and interacting proteins enriched in proteins significantly upregulated in MDMs treated with DDLPS EVs+ cGAMP relative to cGAMP only. (F) Protein-protein interaction network for STING interacting proteins significantly increased in MDM treated with DDLPS EV141 or 246 + cGAMP relative to cGAMP; Visualization represents full STRING network by STRING PPI for high confidence interactions (>0.9) with Markov clustering algorithm inflation parameter 3.

Together, these findings indicate that DDLPS-derived EVs suppress activation of the STING-TBK1-IRF3 signaling axis in macrophages, providing a mechanistic basis for the reduced type I interferon responses and immunosuppressive phenotype observed following EV exposure.

To further characterize the impact of DDLPS EV exposure on cGAMP-induced macrophage programming, we performed Data-Independent Acquisition (DIA) proteomic profiling of MDMs treated with cGAMP alone, or together with EVs isolated from Lipo141 and or Lipo246 cells. Proteomic analysis was performed on cells isolated from three independent donors. Proteomic characterization identified more than 6,000 proteins across all conditions (Figure S4A, B). Consistent with our flow cytometry findings, DDLPS EV-treated macrophages exhibited increased PD-L1 expression together with reduced expression of MHC-II and CD86 relative to macrophages treated with cGAMP alone (Figure 4B). Importantly, EVs derived from both DDLPS cell lines induced highly concordant proteomic changes (Figure 4C).

We next examined proteins significantly altered in both DDLPS EV- and cGAMP-treated MDMs relative to cGAMP only-treated samples: DDLPS EV-treated macrophages exhibited significantly increased expression of 1,036 proteins and reduced expression of 782 proteins, corresponding to 16.9% and 12.7% of the detected proteome, respectively (Figure 4D).

To better define the functional consequences of DDLPS EV-induced changes in macrophage proteome expression, we next performed pathway enrichment and protein-protein interactions analyses. ToppGene analysis(23) and candidate gene prioritization identified significant enrichment of STING1-interacting proteins among proteins increased following DDLPS EV exposure relative to cGAMP treatment alone (Figure 4E). Search tool for the retrieval of interacting genes/proteins (STRING) analysis of STING1-interacting proteins identified enrichment of clusters associated with vesicle biogenesis (clusters 6 and 10) as well as proteasome degradation (cluster 3) and translation (clusters 1 and 4) (Figure 4F). In contrast, analysis of proteins significantly downregulated in DDLPS EV-treated samples identified significant enrichment of genes regulated by class II MHC transactivator (CIITA) protein (Figure S4C), a core positive regulator of MHC-II gene expression. Consistent with these observations, STRING protein-protein interaction analysis from CIITA targeted genes supported attenuated MHC-II antigen processing and presentation of exogenous peptides (cluster 4) in DDLPS EV-treated MDMs (Figure S4D).

Collectively, these findings indicate that DDLPS-derived EVs induce broad macrophage proteome remodeling characterized by attenuation of STING-associated signaling and suppression of MHC-II-dependent antigen presentation pathways.

### EV-associated miRNA signature analysis links DDLPS-derived EVs to immune suppression and clinical outcomes

To place our findings in a clinical context, we first analyzed STING1 expression in DDLPS patients (n=48) from the TCGA sarcoma data set. Relative to patients with lower STING1 expression (n=24), patients with greater than median STING1 mRNA expression (n=24) exhibited prolonged overall survival (Figure 5A). These findings are consistent with an antitumor role for STING signaling in DDLPS.

**Figure 5:**
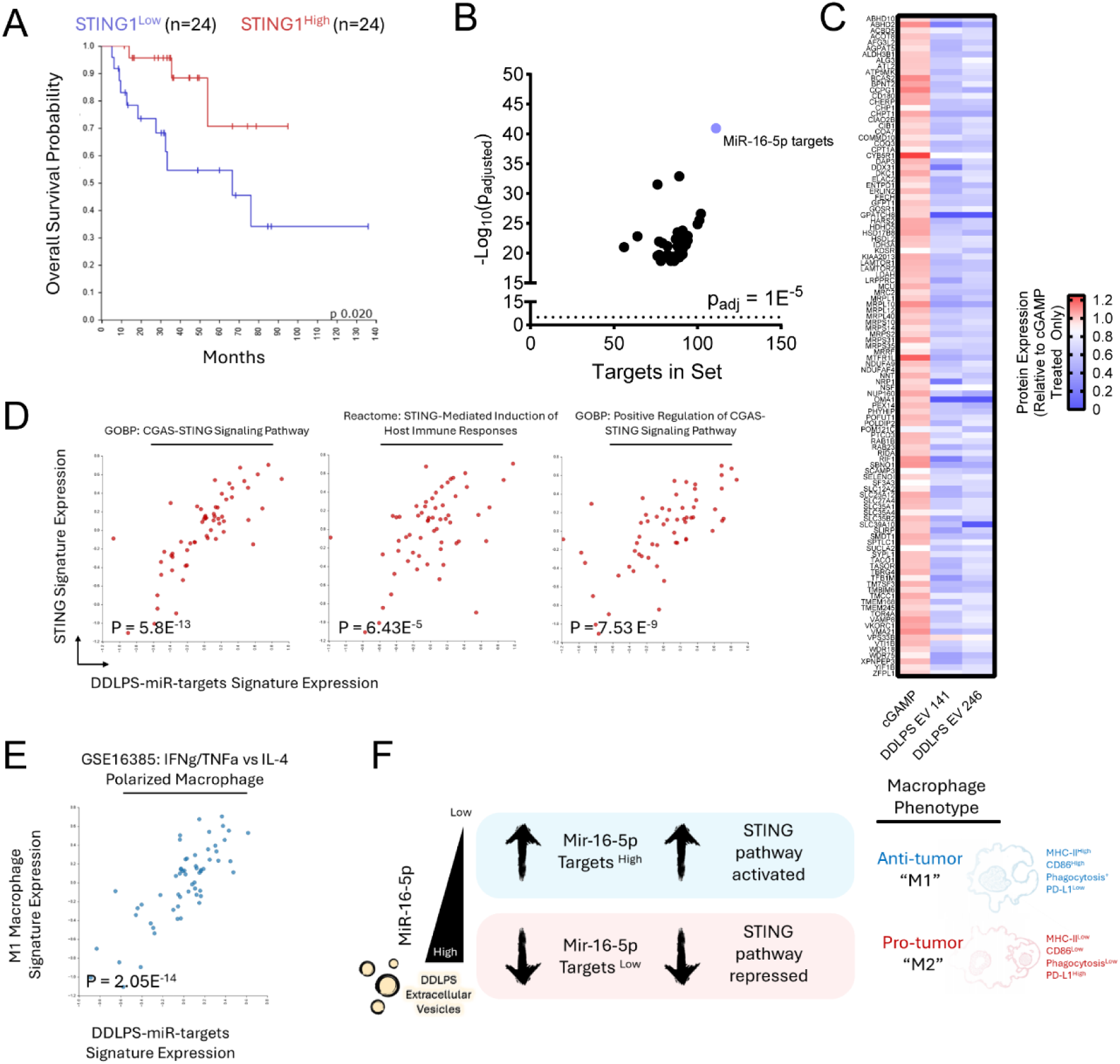
MiR-16-5p targets are negatively coregulated with STING activation in DDLPS: (A) Overall survival comparison for DDLPS patients from the TCGA-Sarcoma cohort split by median expression of STING1 mRNA with STING1 high n=24 and STING1 low n=24. (B) Visualization of miRNA-target pathways enriched in proteins significantly downregulated in MDMs treated with DDLPS EVs+ cGAMP relative to cGAMP only and (C) heatmap visualization of miR-16-5p target proteins downregulated in MDMs treated with DDLPS-EVs. (D) Pathway correlation analysis between miR-16-5p targets downregulated in DDLPS EV treated MDM and STING associated pathways including “GOBP-cGAS-STING-Signaling-Pathway”, GO:0140896, p = 5.8E-13 (*left)* ; “REACTOME-STING-Mediated-Induction-of-Host-Immune-Responses”; R-HAS-1834941, p = 6.43E-5 (*center)*; “GOBP-Positive-Regulation-of-CGAS-STING-Signaling-Pathway”, GO:0141111, p = 7.53E-9 (*right*) and an (E) M1 polarized macrophage signature (“12H-IFNg-TNF-Treated-Macrophage-Up”, GSEA16385, p = 2.05E-14). (F) Schematic representation of proposed relationship between miR-16-5p targeted proteins, STING pathway activation and macrophage phenotypes after DDLPS-EV exposure.

Prior studies from our group identified EV-associated miRNAs as regulators of tumor-associated macrophages in DDLPS(13). Among EV-associated miRNAs previously implicated in DDLPS biology, miR-16-5p is of particular interest because it has been reported to suppress STING signaling in multiple myeloma(24), and we previously demonstrated increased levels of miR-16-5p in plasma-derived EVs from DDLPS patients compared to healthy donors(13). We therefore investigated whether proteins altered in MDMs following DDLPS EV exposure were enriched for miR-16-5p-associated targets.

Notably, pathway analysis of proteins significantly downregulated in DDLPS EV-treated macrophages identified significant enrichment of consensus functional miRNA target interactions for validated miR-16-5p (111 proteins, adjusted p = 1.13E^-42^; “hsa-miR-16-5p: Functional MTI”), hereafter designated as DDLPS-miRNA-targets (Figure 5B). Consistent with a potential contribution of EV-associated miR-16-5p to these changes, expression of these target proteins was broadly reduced in MDMs following DDLPS EV exposure (Figure 5C). Protein-protein interaction analysis for miR-16-5p targets downregulated in DDLPS EV-treated MDMs identified several functionally associated clusters, including mitochondrial translation and gene expression as well as soluble N-ethylmaleimide-sensitive factor attachment protein receptors (SNARE)-mediated vesicular transport pathways (Figure S5A).

We next examined the relationship between DDLPS-miRNA-target expression and STING-associated pathways. Expression of DDLPS-miRNA-targets was strongly associated with multiple signatures linked to STING signaling and pathway activation, including cGAS-STING signaling (GO:0140896, p = 5.8E^-13^), positive regulation of cGAS-STING signaling (GO:0141111, p = 7.53E^-9^), and STING-mediated induction of host immune responses (R-HAS-1834941, p = 6.43E^-5^) (Figure 5D). Moreover, DDLPS-miRNA-target expression was significantly associated with expression of an immunostimulatory macrophage “M1” signature (“12H-IFNg-TNF-Treated-Macrophage-Up”, GSEA16385, p = 2.05E^-14^) (Figure 5E).

Together, these analyses identify a miR-16-5p-associated protein signature positively correlated with STING pathway activation and macrophage M1 polarization. In the context of DDLPS EV exposure, downregulation of miR-16-5p target proteins may therein contribute to reduced STING signaling and impaired macrophage-mediated antitumor immune responses.

## Discussion

In this study, we demonstrate that DDLPS-derived EVs attenuate macrophage responses to cGAMP-mediated STING activation, leading to impaired antitumor immune function (Fig. 6). DDLPS-derived EVs from tumor cell lines and patient serum reduced cGAMP induced STING activation and impaired type I interferon secretion. DDLPS-derived EVs also reprogrammed macrophages towards an immunosuppressive phenotype, impaired phagocytosis, and suppressed macrophage dependent T cell proliferation while increasing T cell exhaustion. Integrated proteomic characterization of DDLPS-EV exposed macrophages confirmed that DDLPS-EV exposure promoted immunoregulatory macrophage polarization. Moreover, DDLPS-EV exposure broadly restructured the proteome of primary human macrophages, including suppression of proteins enriched for validated miR-16-5p targets. Analysis of miR-16-5p targets downregulated after DDLPS-EV exposure indicated a strong association between miR-16-5p target protein expression and STING signaling. These findings suggest that miR-16-5p-associated pathways may contribute to EV-mediated suppression of STING signaling and antitumor macrophage polarization.

**Figure 6:**
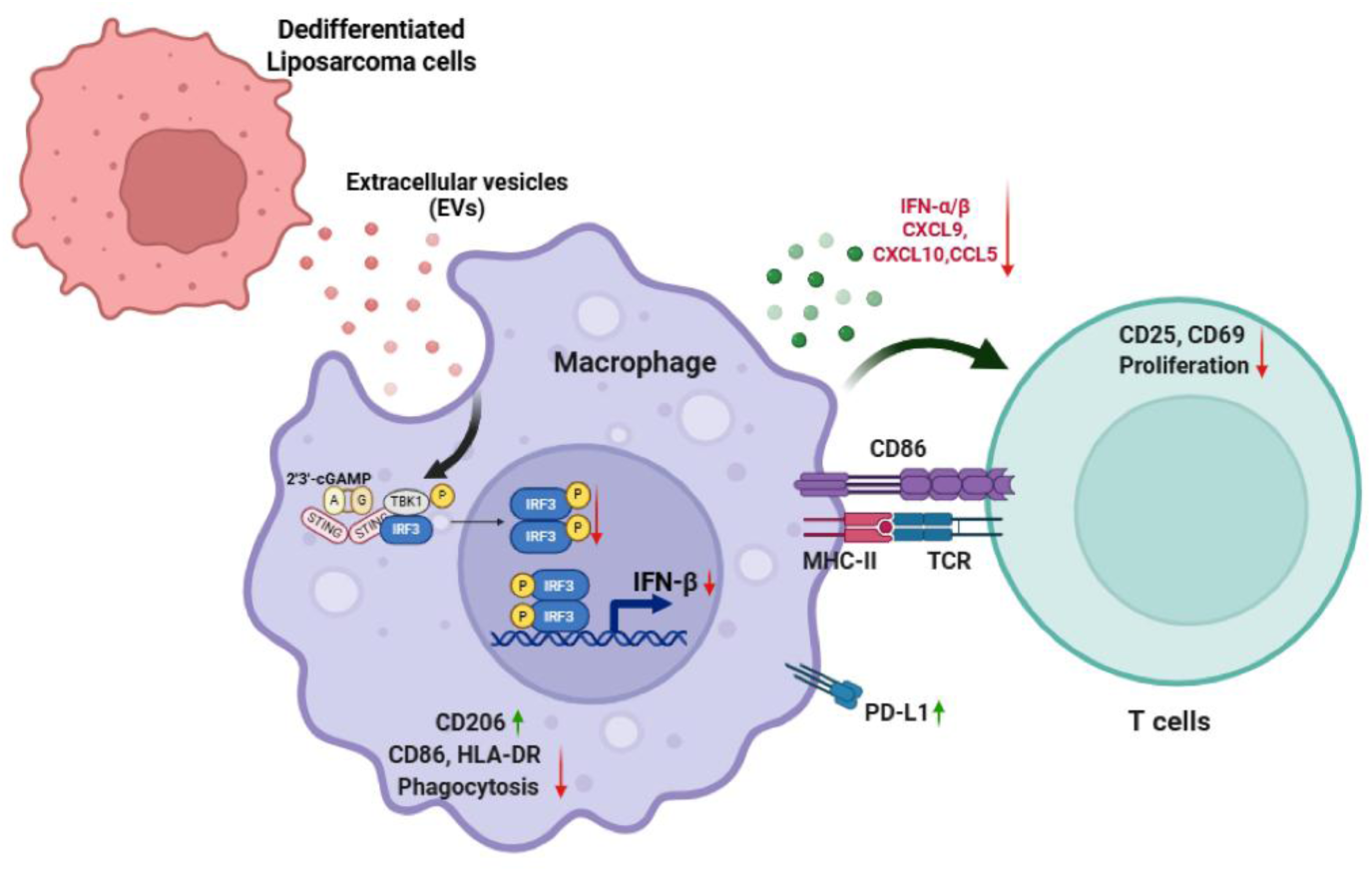
Schematic representation of proposed relationship between DDLPS-EVs, STING signaling, macrophage polarization and T cell responses. DDLPS-derived extracellular vesicles suppress STING signaling in macrophages, thereby attenuating 2,3-cGAMP-induced type I interferon responses, promoting an immunosuppressive macrophage phenotype, and impairing downstream T-cell activation, ultimately contributing to immune evasion in DDLPS.

To our knowledge, this study provides the first evidence that DDLPS-derived EVs suppress STING signaling in macrophages and contribute to impaired antitumor immune responses. Prior reports on the intratumoral immune environment in DDLPS underline a central role for macrophages. Among soft tissue sarcomas, DDLPS exhibits the highest macrophage and CD8+ T cell infiltration scores(25). However, the clinical benefit from immune checkpoint blockade in DDLPS remains relatively modest compared with undifferentiated pleomorphic sarcoma(6, 26). The coexistence of abundant macrophage and T-cell infiltration with limited responsiveness to immunotherapy suggests that immune dysfunction in DDLPS may arise from qualitative defects in immune cell activation rather than a simple absence of immune infiltrates. In clinical trials involving multiple solid tumors, STING agonists have shown limited efficacy in sarcoma compared with other solid tumors(7, 27). This pattern suggests that the defects in antitumor immune activation may represent major barrier to effective immunotherapy in DDLPS. STING signaling in macrophages is particularly important in antitumor immunity because it plays a critical role in dsDNA sensing, regulating IFN-beta production and downstream T cell cytotoxic immune activity(28, 29). A recent study in breast cancer demonstrated that macrophage STING activation can reprogram M2-like tumor associated macrophages toward an M1- like antitumor state, further supporting the importance of STING signaling in macrophage-driven antitumor immunity(18). EV-mediated inhibition of the cGAS-STING pathway in macrophages has also recently been described in glioblastoma(30). TME including macrophage derived chemokines and T cell status are crucial for effective responses to immune checkpoint blockade(31, 32). Therefore, suppression of STING signaling and resulting alterations in macrophage and T-cell function may have important implications for therapeutic responsiveness. In lung cancer, STING suppression has been mechanistically linked to immune escape and resistance to immunotherapy(33). Our findings that DDLPS-derived EVs suppress STING phosphorylation, reduce expression of *IFNB1*, *CXCL9*, *CXCL10*, and impair downstream T cell responses suggest that EV-mediated inhibition of macrophage STING signaling attenuates immune functions required for effective antitumor immunity. This process may partially contribute to the limited efficacy of immunotherapy observed in DDLPS.

Several recent studies connect tumor-derived EV cargos to STING suppression. Tumor exosomal ENPP1 can hydrolyze extracellular cGAMP and thereby inhibit cGAS–STING signaling, providing an important mechanism by which tumor may limit STING activation within the TME(34). In glioblastoma, tumor-derived EVs transfer miR-25/93 into macrophages and impair STING activation, leading to reduced immune response in macrophage and T cells(30). Although the specific cargo responsible for STING suppression in DDLPS remains to be experimentally defined, our data on miRNA signature analysis supports the hypothesis that EV-associated cargo contributes to STING inhibition and identifies potential candidates for future studies.

There are several limitations in this study. First, although both DDLPS cell line-derived EVs and patient serum-derived EVs suppress macrophage interferon responses, the precise EV cargo responsible for STING inhibition remains to be fully defined. The candidate miRNA signature provides a potential starting point, but direct loss of function experiments will be required to determine whether specific EV miRNAs mediate the observed phenotype. Second, most functional experiments were performed *in vitro* using U937-derived macrophages or human monocyte-derived macrophages. Because well-established immunocompetent models of DDLPS remain limited, future studies using humanized mouse systems may help validate these findings *in vivo*.

### Conclusion

DDLPS-derived EVs impair macrophage responsiveness to cGAMP by suppressing STING pathway activation, leading to reduced type I interferon signaling, impaired chemokine expression and T-cell dysfunction. Together, these findings identify DDLPS-derived EVs as regulators of macrophage STING signaling and provide insights into how EV-mediated macrophage reprogramming contributes to impaired antitumor immunity. Targeting the EV-STING axis may therefore represent a potential therapeutic strategy to restore immune responsiveness in DDLPS.

## Materials and Methods

### Patients and clinical samples

Peripheral blood samples from patients with DDLPS and healthy donors were used for isolation of serum-derived EVs. Written informed consent was obtained from all participants, and all human sample collection and use were performed under approval of the institutional review board in accordance with the Declaration of Helsinki. Blood was collected into blood collection tubes, processed to obtain serum used in each experiment, aliquoted, and stored at −80°C until EV isolation.

### Cell lines and culture conditions

Human DDLPS cell lines Lipo141 and Lipo246 were used as EV donor cells. Tumor cells were cultured in DMEM supplemented with 10% fetal bovine serum. For EV collection, cells were transferred to serum-free medium for 48 hours prior to collection of conditioned media. U937 monocytes were cultured in RPMI 1640 supplemented with 10% fetal bovine serum (FBS). To generate macrophage-like cells, U937 cells were differentiated by phorbol 12-myristate 13-acetate (Sigma, 100 ng/mL) in RPMI 1640 Medium (Gibco, # A1049101) without FBS for 24 hours, followed by washing and recovery in complete medium before treatment. Cell lines were confirmed to be free of mycoplasma contamination.

### Isolation and differentiation of primary human macrophages and T cells

Human peripheral blood mononuclear cells were isolated from donor blood by density-gradient centrifugation with Histopaque-1077 (Sigma), and CD14⁺ monocytes were enriched using CD14 microbeads positive selection (Miltenyi Biotec, # 130-050-201). Purified monocytes were differentiated into monocyte-derived macrophages (MDMs) in the presence of 50ng/mL macrophage colony-stimulating factor (PeproTech, # 300-25-10UG) for 7 days, in RPMI 1640 Medium (Gibco, # A1049101) with 10% FBS. T cells were isolated using negative selection by Pan T cell isolation kit (Miltenyi Biotec, # 130-096-535).

### EV isolation

For tumor cell-derived EVs, conditioned medium from Lipo141 and Lipo246 cultures was collected and EVs were isolated by differential ultracentrifugation using the workflow described in our previous publication(13). For serum-derived EVs, vesicles were isolated using ExoQuick (System Biosciences, #EXOQ20A-1) according to the manufacturer’s instructions. EV preparations were resuspended in sterile PBS and quantified before downstream use. Isolated EVs were characterized by transmission electron microscopy (TEM), nanoparticle tracking analysis (NTA), and immunoblotting for canonical EV markers, including CD9 and TSG101, and for the negative marker calnexin. Transmission electron microscopy (TEM) confirmed the presence of vesicular structures consistent with EV morphology (Figure S1A). To characterize EV concentration and, we conducted Nanoparticle Tracking Analysis (NTA) (Figure S1B). Utilizing the same input volume of conditioned medium, we observed comparable EV size distribution from Lipo141 and Lipo246 EVs. To verify EV identity according to Minimal information for studies of extracellular vesicles, we performed western blot and confirmed enrichment of EV markers cluster of differentiation 9 (CD9) and tumor susceptibility gene 101 (TSG101) together with absence of Calnexin (Figure S1C).

### RNA extraction and real-time PCR

Total RNA was extracted from macrophages using RNeasy mini kit (QIAGEN, #74104) according to the manufacturer’s instructions. Complementary DNA was synthesized using High-Capacity cDNA Reverse Transcription Kit (Applied Biosystems, #4374966), and quantitative real-time PCR was performed using Taqman probes with TaqMan Fast Advanced Master Mix (Applied Biosystems, #4444557). Expression of *IFNB1, IFNA2, CCL5, CXCL9, CXCL10*, and *ISG15* was measured in triplicate and normalized to *GAPDH*. Relative expression was calculated using the 2^-ΔΔCt method.

### ELISA

3.5×10^4^ U937cells or MDMs were cultured in a 12-well plate. After the treatment, cell culture supernatants were collected, cleared by centrifugation, and stored at −80°C until analysis. Secreted human IFN-β was quantified using Human IFN-β ELISA Kit (Invitrogen, #414101) according to the manufacturer’s protocol. IFN-β concentrations were calculated from standard curves and adjusted for dilution factors.

### Immunoblot analysis

Whole-cell lysates were prepared in ice-cold Cell Signaling cell lysis buffer supplemented with protease inhibitors and phosphatase inhibitors. Protein concentration was determined by spectrophotometer using Bio-Rad Protein Assay. Protein samples were loaded on 4%–20% precast gels and transferred to nitrocellulose membranes. Membranes were blocked in 5% bovine serum albumin and incubated with primary antibodies against phospho-STING (Cell Signaling, #50907), total STING (Proteintech, # 66680-1-Ig), phospho-TBK1 (Cell Signaling, #5483), total TBK1 (Cell Signaling, #51872), phospho-IRF3 (Cell Signaling, #4947) and total IRF3 (Proteintech, #66670-1-Ig). IRDye 800CW Donkey anti-Rabbit IgG (Licorbio, #926-32213) and IRDye 680RD Donkey anti-Mouse IgG (Licorbio, #926-68072) were used as secondary antibodies. Bands were detected by chemiluminescence using film development or by fluorescence using LI-COR Odyssey CLx.

### Flow cytometry for macrophage phenotype and polarization profiling

For flow cytometric analysis, peripheral blood derived macrophages were harvested by manual detachment from culture plates and washed once with cold DPBS (Ca/Mg free). After washing cells were stained with fixable viability stain (Thermofisher, #L23105), anti-human Fc blocking reagents (BD, #564219), and antibody panels indicated below (Supplementary Table 1). Cells were stained for 30 minutes on ice shielded from light in DPBS (Ca/Mg free) followed by pelleting (1KG x 5min, 4C) and washing once with DPBS (Ca/Mg free) followed by resuspension in DBPS and data acquisition on a 5L Cytek Aurora spectral flow cytometer. Analysis was performed on Flowjo (version 10) or Omiq from live, singlet populations.

### Phagocytosis assay

To compare the phagocytic capacity of peripheral blood derived monocyte/macrophages exposed to extracellular vesicles from DDLPS tumor cells, monocyte/macrophage cultures were harvested after treatment via manual detachment from culture plates and washed once with room temperature 1x Hanks buffered saline solution (Gibco, #14065-056). Cells were resuspended in 1x HBSS warmed to 37C containing fluorescently labeled beads (Sigma, #L3280-1ML) at 10uL/mL and incubated in 200uL/sample at 37C for 20 minutes shielded from light. After incubation, cells were pelleted (1KG x 5min, 4C) and resuspended in 1X HBSS for data acquisition on a 5L Cytek Aurora spectral flow cytometer.

### Coculture of macrophages and T cells

For cocultures, macrophages were treated with DDLPS derived EVs for 24 hours followed by exposure to 10ug/mL 2,3-cGAMP (Invivogen, #tlrl-nacga23-1) for 4 hours. After 4 hours, cGAMP containing media was removed and replaced with complete media for 1 hour to enable removal of residual cGAMP. After 1 hour, media was removed and replaced with fresh media containing 1.7*10^5 peripheral blood derived T cells. For T cell activation assays, T cells were cocultured with monocyte/macrophages for 24 or 48 hours; for T cell proliferation assays, T cells were labeled with Cell trace Violet (Thermofisher, #C34557) and incubated with monocyte/macrophages for 72 hours. For cell trace violet labeling, T cells were counted and resuspended at 4E6 cells/ml with 1uM concentration of cell trace violet for 15min in DPBS (Ca/Mg free) at 37C shielded from light. Excess cell trace violet was quenched by the addition of 5ml complete cRPMI followed by pelleting (1KG x 5min, 4C) and resuspension in fresh complete T cell media for coculture with monocytes/macrophages. For proliferation and activation controls T cells were exposed to dynabeads (Gibco, #11131D).

### Proteomics sample preparation and analysis

After culture with DDLPS EVs and cGAMP, MDMs from three independent donors were harvested via manual detachment, washed three times with PBS and preserved as dry cell pellets in low protein binding tumors at −80C prior to processing. Samples were lysed with proteomics lysis buffer (composed of 50mM tetraethylammonium bromide (TEAB) pH 8.5, 4mM MgCl_2_, 5% ultrapure sodium dodecyl sulfate (SDS) and 1x HALT protease inhibitor in ultrapure MiliQ water) on a Bioruptor sonicator with 15 rounds of 60 seconds sonication and 30 seconds rest at 4C. After sonication, samples were pelleted to remove debris at 15KG for 10 minutes at 14C and supernatant retained. Protein concentration in clarified samples was determined with a Pierce BCA protein assay kit and equivalent amounts of protein used for downstream digestion. Samples were reduced with 20mM dithiothreitol (DTT) at 95C for 10 minutes followed by alkylation with 40mM iodoacetamind (IAA) shielded from light for 1 hour at room temperature. Alkylated samples were quenched with 20mM DTT and incubated for 15 minutes at room temperature shielded from light. Samples were then acidified with 12% phosphoric acid and acidic pH validated by pH strip. Acidified samples were combined with column binding buffer (composed on 90% methanol and 100mM TEAB) and loaded onto Protifi S-Traps (#C002-MINIX-0080PK) followed by washing three times with column binding buffer.

Pierce proteomics grade trypsin was added to S-Trap columns in digestion buffer (composed of 50mM TEAB in MiliQ ultrapure water) at 0.5ug/ml and incubated at 47C for 2 hours. After digestion, samples were eluted with 50mM TEAB (for neutral peptides), 0.2% formic acid (for hydrophilic peptides) and 50% acetonitrile with 0.2% formic acid (for hydrophobic peptides) eluted peptides were pooled and dried via speed vacuum. Dried samples were resolubilized in mobile phase A (composed of 0.1% formic acid in MiliQ water) and sonicated for 10 minutes in an Emerson Branson ultrasonic water bath followed by centrifugation at 13KG for 10minutes to remove insoluble material. Peptides in clarified samples were quantified via nanodrop and diluted to 0.2ug/ul in Mobile phase A in 20ul with addition of 0.5ul iRT spike in (Biognosys).

Samples were acquired on an Orbitrap Exploris 480 mass spectrometer with an inline easy nLC-1200 (Thermofisher) with injection of 2ul/sample. Peptides were separated by an isocratic combination of mobile phase A (composed on 0.1% formic acid in MiliQ water) and mobile phase B (80% acetonitrile with 0.1% formic acid) on an 60cm Aurora column (IonOpticks) with a 1.8kv easy spray emitter. Samples were acquired over 160 minutes per sample at a 250nL flow rate.

For analysis, data was processed via FragPipe-24.0 (MSFragger version 4.4.1) with a Basic search workflow referenced against the human peptide reference (Uniprot, accessed 5-18-2026) with decoy protein inclusion (50%). For peak matching precursor mass tolerance PPM was set to −20 to 20 with mass calibration and parameter optimization and isotope error 0/1/2/3. Protein digestion input was designated as ‘strict trypsin’ with peptide lengths of 7 – 50 and peptide mass range 500 – 3,000Da. Included variable modifications included methionine oxidation and N-terminal acetylation; included fixed modifications included cysteine carbamidomethylation. For validation, PSM validation was performed with Percolator, and protein inference was performed via ProteinProphet. Search results were used to generate a spectral library with b and y ions, with retention time normalized by iRT spike ins and ion mobility calibration performed by automatic selection of a run of reference. Quantification was performed by DIA-NN with false discovery rate set to 0.1 with robust LC for high precision. FragPipe-Analyst was used for visualization and downstream analysis.

The ToppGene tool suit (https://toppgene.cchmc.org/) was used to identify protein interactions, pathways and targes subject to miRNA regulation enriched in proteins up or downregulated in samples treated with DDLPS EVs and cGAMP relative to cGAMP treatment alone; P values represent values with Bonferroni multiple comparison correction. To analyze functional associations of miR-16-5p target proteins downregulated in DDLPS EV treated MDM, protein targets (n=111) identified as “hsa-miR-16-5p: Functional microRNA-target interactions” via ToppGene analysis were compared against consensus annotated genesets via the R2 (r2.amc.nl) geneset versus geneset correlation analysis module. Expression of hsa-miR-16-5p: Functional microRNA-target interactions targets in TCGA DDLPS patient samples were visualized with z-score transformation and a minimal geneset size of 5. For protein-protein interaction analysis the STRING PPI suit (https://string-db.org/) was used with STRING networks restricted to high confidence interactions (>0.9) with Markov clustering algorithm (inflation parameter = 3) and exclusion of unconnected nodes.

### Survival analysis

For survival analysis, DDLPS patients with survival information and transcriptomic data from the TCGA Sarcoma (TCGA, Firehose Legacy) cohort were stratified by median mRNA expression of STING1 resulting in n=24 for STING1 high patients and n=24 for STING1 low patients. Overall survival was compared via Gehan-Breslow-Wilcoxon test comparison.

### Statistical analysis

Data is presented as mean ± SEM. Statistical analyses were performed using GraphPad Prism 10.6.0. Comparisons between two groups were evaluated using two-tailed unpaired Student’s t tests, whereas comparisons among multiple groups were performed using one-way ANOVA followed by Dunnett’s multiple-comparisons test. For statistical comparisons *p*< 0.05 corresponds to *, *p*< 0.01 **, *p*< 0.001 *** and *p*< 0.0001 ****.

## Acknowledgements

We thank the Genomics Shared Resource at The Ohio State University Comprehensive Cancer Center, Columbus, OH, for the support with qRT-PCR analysis.

## Informed Consent Statement

Informed consent was obtained from all subjects involved in the study.

