## Supplemental Figure 1-5 for "Extracellular vesicle-mediated suppression of macrophage STING signaling promotes immune dysfunction in dedifferentiated liposarcoma"

**Supplemental Figures and Figure Legends:**

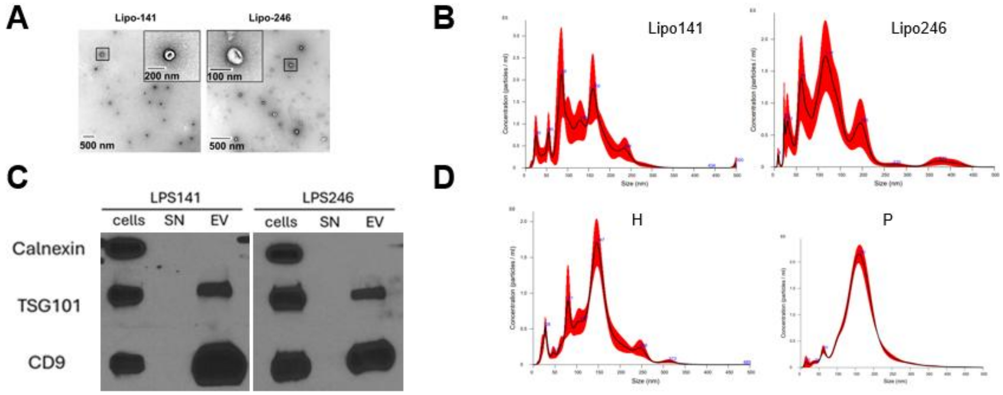


**Figure S1:** (A) Representative transmission electron microscope images of EVs isolated from Lipo141 and Lipo246 cell lines. (B) Representative histograms of extracellular vesicles derived from Lipo141, Lipo246 cell lines. Axis X = size distribution [nm]. Axis Y = concentration [$10^5$ particles/ml].(C) Western blot analysis for Calnexin, CD9 and TSG101 of cells lysates, supernatant (SN) and EVs isolated from Lipo141 and Lipo246 cell lines. (D) Representative histograms of extracellular vesicles derived from serum of healthy donors (H) and DDLPS patients (P). Axis X = size distribution [nm]. Axis Y = concentration [$10^6$ particles/ml].

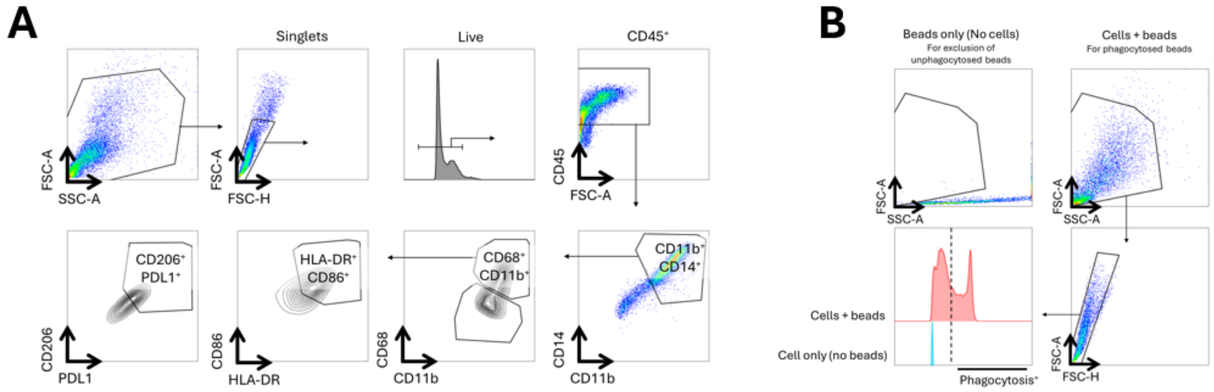


**Figure S2:** (A) Flow cytometric gating approach for identification of macrophage phenotypes and (B) phagocytosis from data represented in Figure 2.

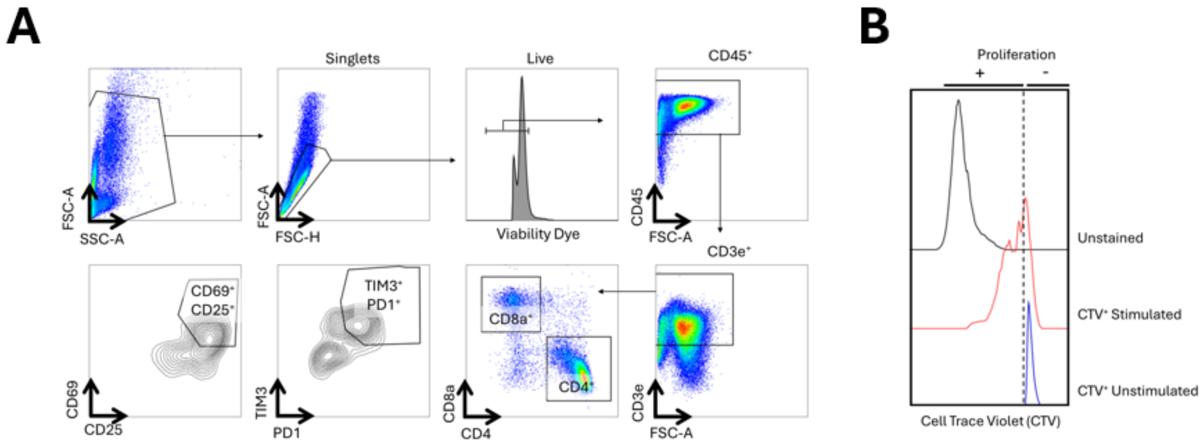


**Figure S3:** (A) Flow cytometric gating approach for identification of T cell phenotypes and (B) proliferation after coculture with macrophages from data represented in Figure 3.

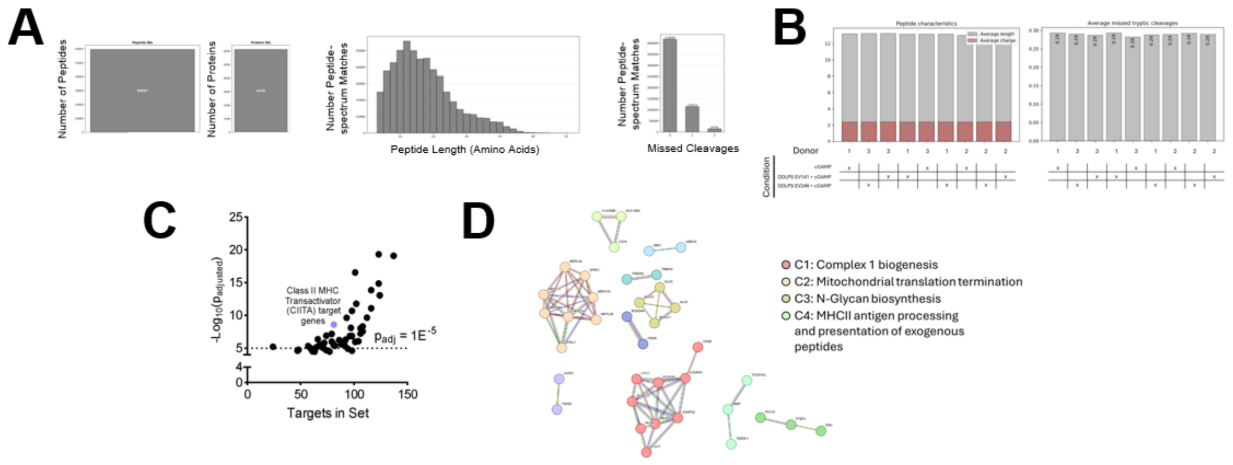


**Figure S4:** (A): Number of peptides and Left: proteins identified across all samples for DIA proteomic characterization of human monocyte derived macrophages exposed to cGAMP or DDLPS EVs and cGAMP (*left*); Number of peptide spectrum matches versus peptide length from proteomic analysis (*center*); Number of missed cleavages across peptide spectrum matches (0 missed cleavages = 76%; 1 missed cleavage = 21%; 2 missed cleavage = 3%)(*right*). (B) Comparison of peptide characteristics for average peptide length and charge across all samples (*left*) and average number of missed tryptic cleavages across all samples (*right*). (C) Visualization of transcription factor binding targets enriched in proteins significantly downregulated in MDM treated with DDLPS EVs+ cGAMP relative to cGAMP only. (D) Protein-protein interaction network for CIITA target genes significantly decreased in MDM treated with DDLPS EV141 or 246 + cGAMP relative to cGAMP; Visualization represents full STRING network by STRING PPI for high confidence interactions (>0.9) with MCL clustering inflation parameter 3

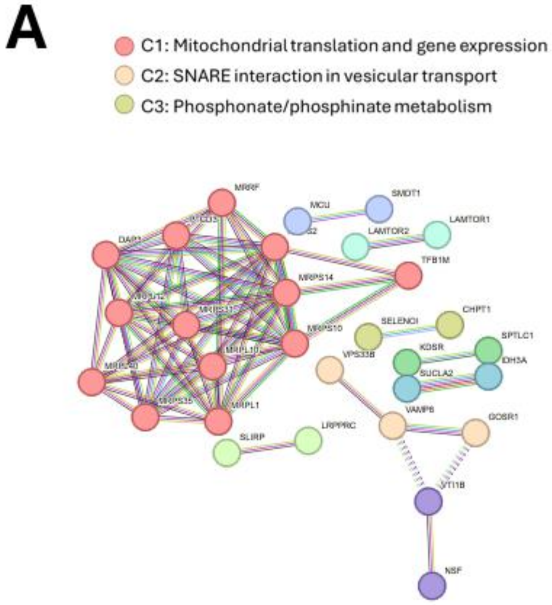


**Figure S5:** (A) STRING protein-protein interaction network for MiR-16-5p targets significantly downregulated in DDLPS-EV exposed MDMs (PPI enrichment $p<1.0E^{-16}$)
